# *Acinetobacter baumannii* sampled from cattle and pigs represent novel clones

**DOI:** 10.1101/2022.03.29.486289

**Authors:** Valeria Mateo-Estrada, Leila Vali, Ahmed Hamouda, Benjamin A. Evans, Santiago Castillo-Ramírez

## Abstract

*Acinetobacter baumannii* is a very important human pathogen. Nonetheless, we know very little about non-human isolates of *A. baumannii*. Here we determine the genomic identity of 15 cattle and pig isolates, as well their antibiotic and virulence genetic determinants, and compare them to the main human clinical international clones.

## Main text

*Acinetobacter baumannii* is a Gram negative opportunistic bacterial pathogen, notorious for being associated with high morbidity and mortality due to its highly drug-resistant nature. While *A. baumannii* can be isolated from clinical samples, its natural home is less clear. Animals have been suggested as a potential host or reservoir for *A. baumannii*. Birds, in particular White Storks, have been proposed as a reservoir [1], though this doesn’t seem to apply to other bird species [2], and as *A. baumannii* can be released into the environment from hospital effluent [3] it isn’t clear the degree to which wild animals are acquiring the bacteria from contaminated soil and water. However, it is clear that *A. baumannii* should be considered a One Health issue, as some non-human isolates have important antibiotic resistance genes [4]. *A. baumannii* seems to be fairly common in domestic livestock, particularly cattle [5, 6], where isolates tend to have a generally susceptible antibiotic resistance profile and appear to be genetically distinct from clinical strains by molecular typing methods. In a previous study, 16 *A. baumannii* isolates were collected from cattle and pigs that had been recently slaughtered, and were shown by PFGE to cluster separately from the 3 major clonal lineages of *A. baumannii* prevalent at the time; furthermore, they carried different *oxaAb* (*bla*_OXA-51-like_) variants [7]. Here, we sequenced the genomes of these 16 isolates to determine how genetically similar they are to human clinical isolates.

Total DNA was extracted from overnight broth cultures with a Promega Wizard Genomic DNA Purification kit (Promega, UK), quality checked by nanodrop and quantity assessed by Qubit. Purified DNA was paired-end sequenced on an Illumina platform. The sequences were trimmed with Trim Galore [8] and assembled via SPAdes [9], as described previously [10]. The genomes were annotated employing Prokka [11] and genotyped by the Pasteur Multilocus Sequencing Typing (MLST) scheme [12] using the PubMLST online database [13]. The genome quality was assessed with CheckM [14] and only the genomes with more than 95% completeness and less than 5% contamination were considered for downstream analyses. One isolate from a pig faecal sample (PF33) was discarded as it showed a high percentage of contamination (>60%). For the phylogenetic analysis, we also included 148 human-related *A. baumannii* genomes previously genotyped in [15]. These genomes were chosen as they are part of the eight main international clones (ICs) [16-18]. A Maximum Likelihood (ML) core phylogeny was built using the strategy described in [19]. Briefly, the genes present in a single copy in all the genomes (single-gene families) were identified with Roary [20] and tested for recombination using PhiPack [21]. We found 759 single-gene families without recombination, which were concatenated to build a phylogeny with RAxML [22] and the tree was annotated with iTOL [23]. The antibiotic resistance gene prediction was carried out with the Comprehensive Antibiotic Resistance Database (CARD) [24], and *ampC* alleles were identified using the PubMLST database.

A ML core genome phylogeny of the 15 animal isolates alongside a collection of 148 clinical isolates representing the major international clones [15] showed that the animal isolates formed 3 well separated clades, each of which was distinct and very distant from any of the clinical isolates (Figure 1). The pig faecal isolates formed a single clade, two of the cattle faecal isolates (CF233 and CF234) formed a second clade, and the remaining 4 cattle faecal isolates formed a clade with the two cattle nostril isolates. Considering the Pasteur MLST genotyping, these three clades corresponded to sequence type (ST^PAS^)162, ST^PAS^1014 and ST^PAS^492 respectively. As described previously, the isolates belonging to ST^PAS^1014 carried the *oxaAb* variant *oxaAb*(150), and the ST^PAS^492 isolates carried *oxaAb*(148) [7]. However, the STPAS162 strains carried *oxaAb*(51), which is considered diagnostic of International Clone (IC)4 isolates [25]. IC4 isolates typically belong to ST^PAS^15, which only shares a single allele in common with ST^PAS^162, and the ST^PAS^162 and IC4 isolates are very clearly separated in the core gene phylogeny (Figure 1). It is interesting to note that incongruence between MLST ST and *oxaAb* allele has been observed previously for ST^PAS^162, and warrants further investigation [26]. Of note is that all published ST^PAS^162 isolates, and all of those in the PubMLST database, are from South American countries (Brazil and Chile), geographically very distant from the Scottish isolates reported here. Only two STPAS492 isolates are listed in the PubMLST database, from Lebanon and Russia, and one of these is an animal isolate. There are no other ST^PAS^1014 isolates in the PubMLST database, but there are 11 isolates that match 6 loci, and of these, two are listed as coming from animals, two from food, and one from the environment, suggesting that these STs are commonly isolated from non-clinical sources. Collectively, these results show that the pig and cattle isolates form well differentiated groups and they are not closely related to the major international clones.

**Figure 1:**
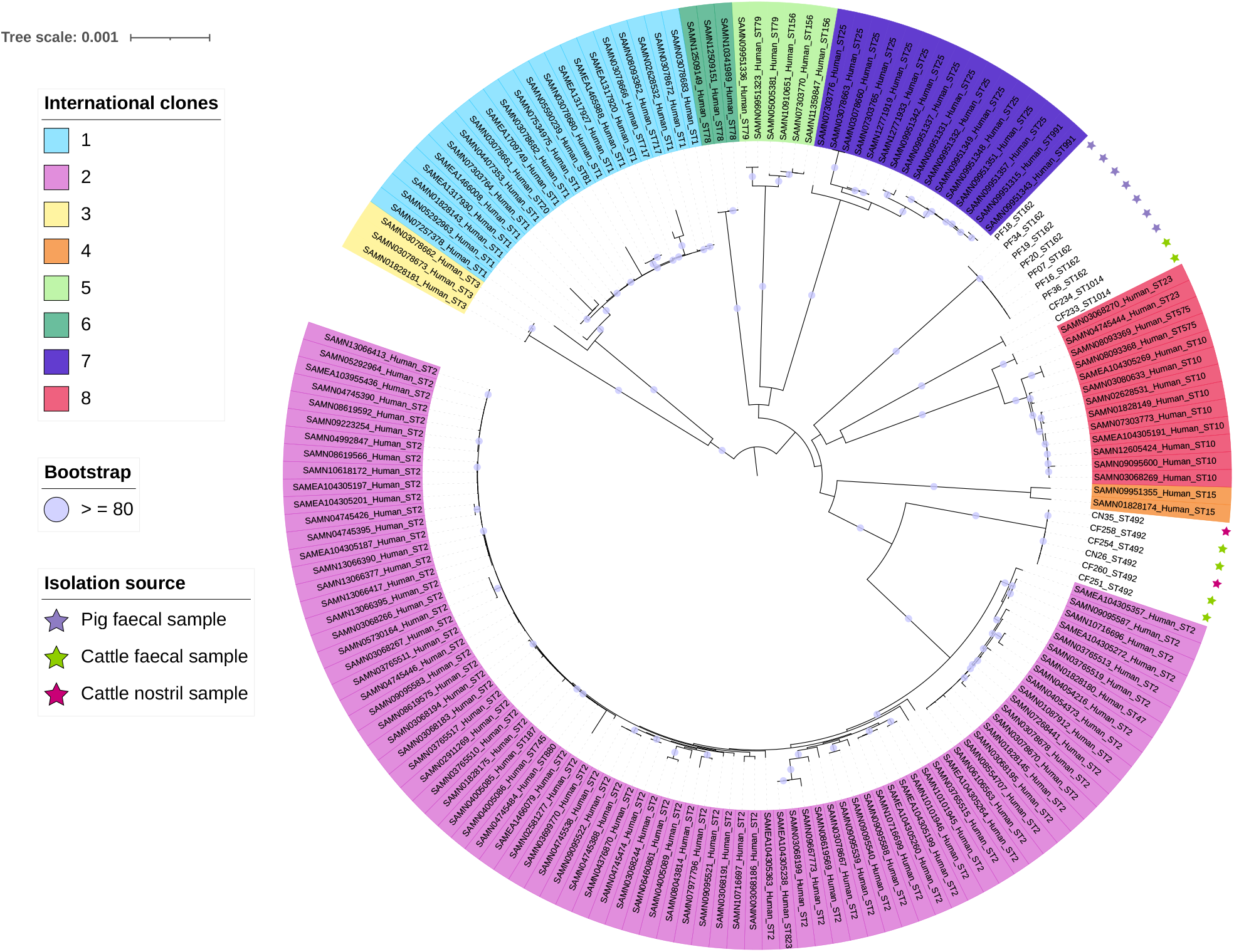
Core genome Maximum Likelihood phylogeny of animal isolates and 148 clinical isolates from 8 international clonal lineages. The major international clones are highlighted with different colours on the labels. Pig isolates are denoted with violet stars, whereas cattle isolates are shown with green (faecal samples) and rosy (nostril samples) stars. The tree scale is in number of substitutions per site and bootstrap values higher (or equal to) 80 are depicted with violet circles at the internal nodes of the phylogeny.

As expected from the previously reported generally antibiotic sensitive nature of the isolates, only chromosomally-encoded resistance genes such as *oxaAb, ampC* and efflux systems were identified, with no acquired antibiotic resistance genes present (Supplementary Figure 1). It had previously been described for these animal isolates that they did not carry IS*Aba1* upstream of the *oxaAb* genes or the *ampC* genes, where it can provide a promotor for their expression [7]. Of the 15 isolates, 7 had no substantial match to any insertion sequences in the ISFinder database (accessed 09/03/2022). Of the remaining 8, 1 isolate (CN26) had a short contig with a partial match to IS*1411*, while all 7 pig faecal isolates contained a 184 bp fragment with 97% identity to the 5’ end of IS*Ajo2*, and a 279 bp fragment with 84% similarity to the 3’ end of IS*Acsp2*. We therefore did not detect any complete IS elements in these strains. In *A. baumannii*, IS elements are thought to be a major mechanism through which the bacteria regulate gene expression and mobilises genes, and are a common feature of clinical isolates. Their almost complete absence from these animal isolates highlights how different in nature they are from clinical isolates, and that IS-mediated adaptation may be a feature of successful clinical strains rather than a general characteristic of the species.

In order to assess whether the animal isolates differed in their complement of virulence factors, they were analysed using VFanalyser alongside 14 clinical and 1 human louse isolate included in the database [27]. Animal isolates had a significantly smaller complement of capsule-related genes, averaging 17, whereas the clinical isolates averaged 22 (t-test, *p* = 0.000016). The six ST^PAS^492 isolates differed from the other animal isolates in that they lacked 8 genes involved in heme utlisation, including *hemO* (Supplementary Table 1). Variation in the carriage of these genes is common, with seven out of the 14 clinical and 1 louse strains also lacking these genes. Thus, these data suggest that the animal isolates may have fewer virulence factors than human clinical isolates.

In conclusion, our study shows that these cattle and pig isolates represent 3 novel lineages well separated from the major international clones. Furthermore, these new lineages are distinct in nature considering both antibiotic resistance and virulence when compared with their human clinical counterparts.

## Funding

This work was supported by CONACyT Ciencia Básica 2016 (grant no. 284276) and “Programa de Apoyo a Proyectos de Investigación e Innovación Tecnológica PAPIIT” (grant no. IN206019) given to SCR. VME is a doctoral student from the Programa de Doctorado en Ciencias Biomèdicas, Universidad Nacional Autónoma de México, and she is funded by a CONACYT doctoral fellowship (no. 1005234).

## Conflict of Interest

The authors declare no conflicts of interest.

## Acknowledgements

We are thankful Ana Mateus from MRCVS, Clinical Scholar, Animal Production and Public Health Department, Glasgow University Veterinary School for supplying the abattoir samples.

## FIGURE LEGENDS

**Supplementary Figure 1:** Matrix showing the presence/absence of genes involved in antibiotic resistance in the animal isolates. The genes are colour-coded according to the drug classes they confer resistance to.

**Supplementary Table 1:** Summary output from VFanalyser of the genes involved in virulence in the animal isolates

## Notes

### Competing Interest Statement

The authors have declared no competing interest.

## References

1. Wilharm G, Skiebe E, Higgins PG, et al. Relatedness of wildlife and livestock avian isolates of the nosocomial pathogen Acinetobacter baumannii to lineages spread in hospitals worldwide. Environ Microbiol 2017; 19(10): 4349–64.

2. Łopińska A, Indykiewicz P, Skiebe E, et al. Low occurrence of Acinetobacter baumannii in gulls and songbirds. Pol J Microbiol 2020; 69: 1–6.

3. Seruga Music M, Hrenovic J, Goic-Barisic I, Hunjak B, Skoric D, Ivankovic T. Emission of extensively-drug-resistant Acinetobacter baumannii from hospital settings to the natural environment. J Hosp Infect 2017; 96(4): 323–7.

4. Hernández-González IL, Castillo-Ramírez S. Antibiotic-resistant Acinetobacter baumannii is a One Health problem. The Lancet Microbe 2020; 1(7): E279.

5. Rafei R, Hamze M, Pailhoriès H, et al. Extrahuman epidemiology of Acinetobacter baumannii in Lebanon. Appl Environ Microbiol 2015; 81(7): 2359–67.

6. Klotz P, Higgins PG, Schaubmar AR, et al. Seasonal Occurrence and Carbapenem Susceptibility of Bovine Acinetobacter baumannii in Germany. Front Microbiol 2019; 10: 272.

7. Hamouda A, Findlay J, Al Hassan L, Amyes SG. Epidemiology of Acinetobacter baumannii of animal origin. Int J Antimicrob Agents 2011; 38(4): 314–8.

8. Krueger F. Trim Galore. https://github.com/FelixKrueger/TrimGalore.

9. Bankevich A, Nurk S, Antipov D, et al. SPAdes: a new genome assembly algorithm and its applications to single-cell sequencing. J Comput Biol 2012; 19(5): 455–77.

10. Mateo-Estrada V, Fernández-Vázquez JL, Moreno-Manjón J, et al. Accessory Genomic Epidemiology of Cocirculating Acinetobacter baumannii Clones. mSystems 2021; 6(4): e0062621.

11. Seemann T. Prokka: rapid prokaryotic genome annotation. Bioinformatics 2014; 30(14): 2068–9.

12. Diancourt L, Passet V, Nemec A, Dijkshoorn L, Brisse S. The population structure of Acinetobacter baumannii: expanding multiresistant clones from an ancestral susceptible genetic pool. PLoS One 2010; 5(4): e10034.

13. Jolley KA, Bray JE, Maiden MCJ. Open-access bacterial population genomics: BIGSdb software, the http://PubMLST.org website and their applications. Wellcome Open Res 2018; 3: 124.

14. Parks DH, Imelfort M, Skennerton CT, Hugenholtz P, Tyson GW. CheckM: assessing the quality of microbial genomes recovered from isolates, single cells, and metagenomes. Genome Res 2015; 25(7): 1043–55.

15. Hernández-González IL, Mateo-Estrada V, Castillo-Ramirez S. The promiscuous and highly mobile resistome of Acinetobacter baumannii. Microb Genom 2022; 8(1).

16. Tomaschek F, Higgins PG, Stefanik D, Wisplinghoff H, Seifert H. Head-to-Head Comparison of Two Multi-Locus Sequence Typing (MLST) Schemes for Characterization of Acinetobacter baumannii Outbreak and Sporadic Isolates. PLoS One 2016; 11(4): e0153014.

17. Levy-Blitchtein S, Roca I, Plasencia-Rebata S, et al. Emergence and spread of carbapenem-resistant Acinetobacter baumannii international clones II and III in Lima, Peru. Emerg Microbes Infect 2018; 7(1): 119.

18. Fonseca É, Caldart RV, Freitas FS, et al. Emergence of extensively drugresistant international clone IC-6 Acinetobacter baumannii carrying blaOXA-72 and blaCTX-M-115 in the Brazilian Amazon region. J Glob Antimicrob Resist 2020; 20: 18–21.

19. Graña-Miraglia L, Evans BA, López-Jácome LE, et al. Origin of OXA-23 Variant OXA-239 from a Recently Emerged Lineage of Acinetobacter baumannii International Clone V. mSphere 2020; 5(1).

20. Page AJ, Cummins CA, Hunt M, et al. Roary: rapid large-scale prokaryote pan genome analysis. Bioinformatics 2015; 31(22): 3691–3.

21. Bruen TC, Philippe H, Bryant D. A simple and robust statistical test for detecting the presence of recombination. Genetics 2006; 172(4): 2665–81.

22. Stamatakis A. RAxML version 8: a tool for phylogenetic analysis and post-analysis of large phylogenies. Bioinformatics 2014; 30(9): 1312–3.

23. Letunic I, Bork P. Interactive Tree Of Life (iTOL) v4: recent updates and new developments. Nucleic Acids Res 2019; 47(W1): W256–W9.

24. Alcock BP, Raphenya AR, Lau TTY, et al. CARD 2020: antibiotic resistome surveillance with the comprehensive antibiotic resistance database. Nucleic Acids Res 2020; 48(D1): D517–D25.

25. Zander E, Nemec A, Seifert H, Higgins PG. Association between β-lactamase-encoding bla(OXA-51) variants and DiversiLab rep-PCR-based typing of Acinetobacter baumannii isolates. J Clin Microbiol 2012; 50(6): 1900–4.

26. Opazo-Capurro A, San Martín I, Quezada-Aguiluz M, et al. Evolutionary dynamics of carbapenem-resistant Acinetobacter baumannii circulating in Chilean hospitals. Infect Genet Evol 2019; 73: 93–7.

27. Liu B, Zheng D, Jin Q, Chen L, Yang J. VFDB 2019: a comparative pathogenomic platform with an interactive web interface. Nucleic Acids Res 2019; 47(D1): D687–D92.

