## Supplementary figures and images for "*Acinetobacter baumannii* sampled from cattle and pigs represent novel clones"

### Supplementary Figure 1

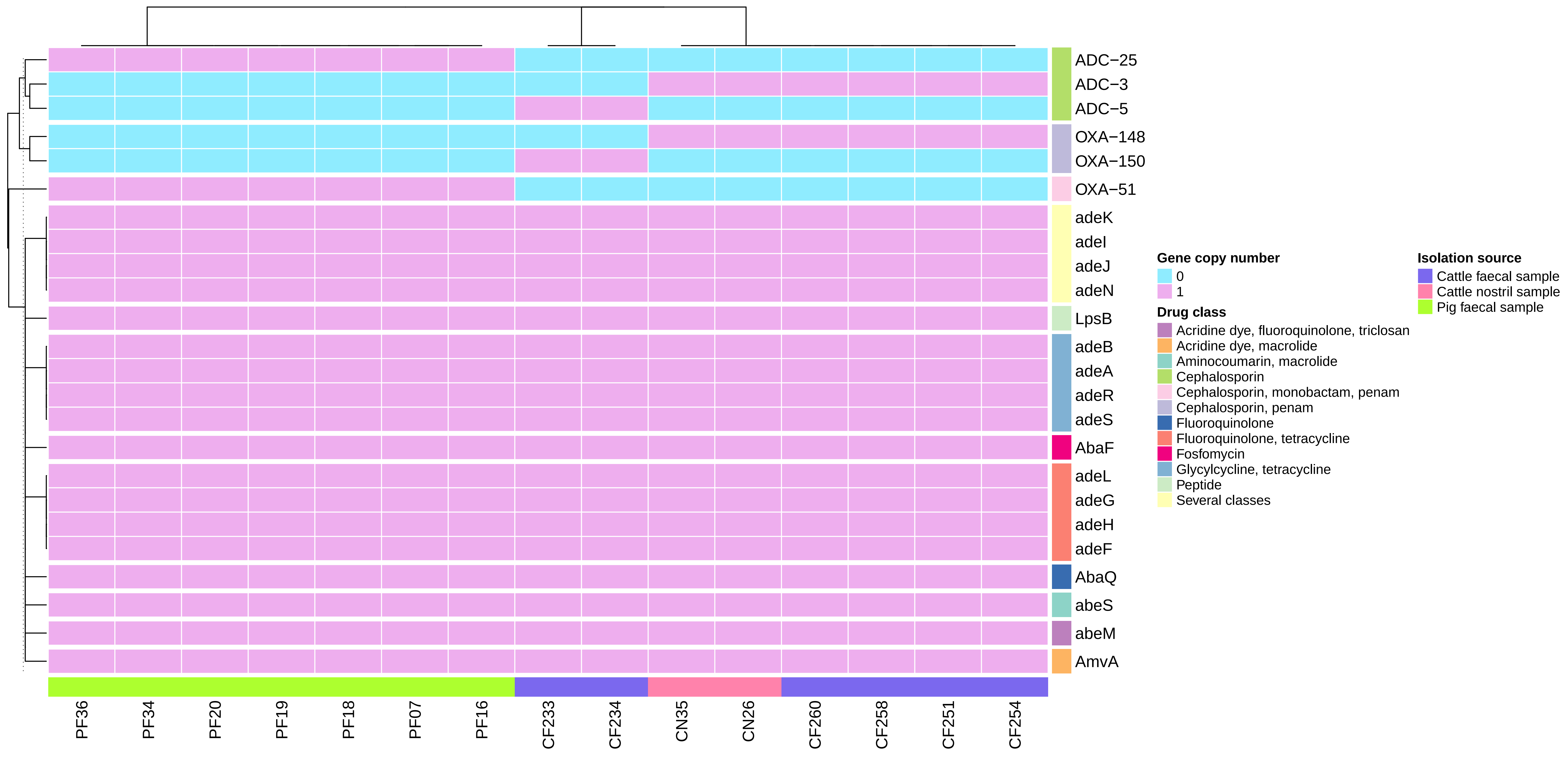
